# Successful Liver transduction by Re-administration of Different Adeno-Associated Virus Vector Serotypes in Mice

**DOI:** 10.1101/2022.10.21.513281

**Authors:** Nemekhbayar Baatartsogt, Yuji Kashiwakura, Takafumi Hiramoto, Morisada Hayakawa, Nobuhiko Kamoshita, Tsukasa Ohmori

## Abstract

Intravenous administration of adeno-associated virus (AAV) vector is a promising gene therapy approach for monogenic diseases. However, re-administration of the same AAV serotype is impossible due to the induction of anti-AAV neutralizing antibodies (NAbs). Here we examined the feasibility of re-administration of AAV vectors to change the serotypes. We administered AAV3B, AAV5, or AAV8 vectors targeting the liver of C57BL/6 mice intravenously, and then assessed the emergence of NAbs and the transduction efficacy with a second administration. For all serotypes, we confirmed that re-administration with the same serotype was not possible. Although the highest neutralizing activity of NAb was induced by AAV5; however, the NAbs elicited by AAV5 did not react with any other serotypes, resulting in success in re-administration with the other serotypes. The re-administration of AAV5 was also successful in all mice treated with AAV3B and AAV8. The effective secondary administration of AAV3B and AAV8 was observed in most mice treated with AAV8 and AAV3B, respectively. However, few mice developed NAbs cross-reactive with the other serotypes, especially the serotypes with close sequence homology. In summary, AAV vector administration induced NAbs relatively specific to the serotype administrated. Secondary administration of AAVs targeting liver transduction could be successfully achieved by switching AAV serotypes in mice.

## Introduction

Gene therapy with an adeno-associated viral (AAV) vector has been attracting attention as a therapeutic gene-delivery vehicle for a variety of inherited monogenic disorders, such as hemophilia.^1–4^ Systemic intravenous injection of liver-tropic AAV vector can efficiently transduce liver hepatocytes to express the target proteins. Recent success in preclinical and human clinical trials of AAV-mediated gene therapy for hemophilia has shown the durable transgene expression at the therapeutic level which dramatically reduced the use of therapeutic coagulation factor concentrates.^5–8^

Although AAV-mediated gene therapy has become an attractive alternative treatment option for hemophilia, there are several compatibility issues to be addressed in order to be useful for many patients. One of the main obstacles in achieving AAV-mediated gene therapy is the presence of circulating neutralizing antibody (NAb) against AAV capsid related to previous AAV infection in some individuals. The therapeutic transduction by systemic administration of an AAV vector is hindered by the NAbs against the AAV, even at low levels.^9–11^ Hence, most clinical trials of systemic administration of AAV vector have targeted only patients without NAbs against the AAV.^12–14^ We have recently reported the prevalence of NAbs against several AAV serotypes in Japanese hemophiliacs and healthy individuals and found the prevalence to range from 20% to 30% in the parent population.^15^

The administration of AAV vectors, in addition to natural infection with AAV, correspondingly induces a high titer of NAbs against the AAV vector. The induction of NAbs against AAV makes it impossible to re-administer AAV vector with the same serotype. Because AAV vectors are not readily inserted into chromosomal DNA, their therapeutic effect is diluted due to cell proliferation. It is possible that the therapeutic effects gradually diminished after further long-term observation. In addition, the therapeutic effect of AAV varies widely among individuals, leading to inadequate efficacy in some patients. Hence, to resolve these inhibitory circumstances, it is necessary to develop a strategy to re-administer the AAV vector.

One of the most practical ways to re-administer the vector is by changing the AAV serotype in the second challenge. The re-administration of the AAV1 vector reportedly succeeded in AAV5-treated monkeys and mice.^16^ In addition, we have reported an increase in plasma coagulation factor IX (FIX) by re-administration of AAV3 vector at five years after its first administration in a cynomolgus monkey previously treated with AAV8.^17^ However, little is known about the reactivity of NAbs induced by AAV vector administration to other serotypes or which AAV serotypes can be used for re-administration. Here, we examined the cross-reactivity of NAbs induced by three different AAV vectors with different serotypes, as well as the possibility of re-administering each of the vectors.

## Materials and Methods

### Comparison of capsid sequence among AAV serotypes

Phylogenetic tree of the capsid sequences (VP1) of AAV serotypes was constructed and aligned with MUSCLE in MEGA 11 software. ^18^ The percentage of identity and similarities was analyzed by Snapgene 5.15 software (GSL Biotech, San Diego, CA).

### AAV vector production

As we have previously described, we produced various serotypes of AAV vector harboring the luciferase gene or the secreted type of the NanoLuc gene driven by the CAG promoter for the determination of anti-AAV NAbs.^15^ For *in vivo* transduction experiments, we created AAV3B, AAV5, and AAV8 vectors harboring human coagulation factor IX variant R338L (FIX Padua) or the secreting type of NanoLuc (secNanoLuc) genes under the control of human α1-antitrypsin promoter conjugated with ApoE hepatic control region (HCRhAAT promoter). The AAV vector was produced by the transfection of AAVpro293 cells (Takara Bio, Shiga, Japan) with three plasmid transfections (Helper-free system). Subsequently, the AAV vector was isolated by CsCl-based ultracentrifugation and the vector titer was determined, as described previously.^19^

### Animal experiments

All animal experiments were approved by the Institutional Animal Care and Concern Committee of Jichi Medical University and were conducted in accordance with the committee’s guidelines and ARRIVE guidelines (permission number #20023-01). C57BL6/J male mice were obtained from Japan SLC (Shizuoka, Japan). We obtained blood from the right jugular vein at an indicated time and intravenously administered AAV vectors into the mouse anesthetized with isoflurane. To obtain liver tissue, mice were deeply anesthetized with isoflurane and then intracardially perfused with phosphate-buffered saline to remove the blood. Then, the liver tissues were dissected for the analysis of AAV vector copy number, as described below.

### Anti-AAV NAb assay

NAb screening and titer were essentially evaluated using a cell-based assay, as described previously.^15^ Briefly, the sera from mice were heat-inactivated (56°C, 30 min) and diluted with FBS, and then incubated with an AAV vector at 37°C for 60 min. The mixture was added into a 96 well plate seeded with Huh-7 cells (Riken BRC, Ibaraki, Japan) at multiplicity of infection of 1,500. The luciferase activity in cell lysate and secNanoLuc activity in cell supernatant were determined by a luminometer (Centro LB 960, BERTHOLD Technologies, Bad Wildbad, Germany). The serum that inhibited >50% vector transduction at 1:1 dilution was considered a positive sample. The sera of positive samples were serially diluted with FBS, and then the inhibition of vector transduction was assessed. We estimated antibody titers that neutralized 50% vector transduction (ND50) by nonlinear regression using GraphPad Prism (GraphPad, San Diego, CA).

### AAV vector copy number in liver

Genomic DNA was extracted from the liver tissue using a nucleic acid extraction system GENE PREP STAR 480 (Kurabo, Osaka, Japan). AAV vector DNA was quantified by qPCR time using THUNDERBIRD Probe qPCR Mix (Toyobo, Osaka, Japan) on QuantStudio 12K Flex (ThermoFisher Scientific). The primer pairs and probe used for the detection of AAV genome were as follows: 5’-AGCAATAGCATCACAAATTTCACAA-3’ (sense) 5’-CCAGACATGATAAGATACATTGATGAGTT-3’ (anti-sense) 5’-AGCATTTTTTTCACTGCATTCTAGTTGTGGTTTGTC-3’ (FAM probe) for the detection of SV40 polyA. 5’-GGAGGTGTGTCCAGTTTGTT-3’ (sense) 5’-ATGTCGATCTTCAGCCCATTT-3’ (anti-sense), and 5’-ATCCCAAGGATTGTCCTGAGCGGT-3’ (FAM probe) for the detection of secNanoLuc.

### Statistical analysis

All statistical analyses were performed using GraphPad Prism (GraphPad, CA, USA). Student’s *t*-test was used to compare two groups. One-way or two-way analysis of variance (ANOVA) and multiple comparisons test were used for calculations that involved more than two-group comparisons. *P*-value of <0.05 was considered statistically significant.

## Results

### Comparison of the capsid amino acid sequence among AAV serotypes

We compared the capsid amino acid sequence (VP1) among several AAV serotypes (Figure 1). To analyze the specificity of NAb after AAV vector was administered, we selected three different clades of AAVs, *e.g*, AAV3B, AAV5, and AAV8 (Figure 1). AAV5 and AAV8, which have been widely used in gene therapy with the liver as the target in clinical trials, and AAV3B, which reportedly possesses a high gene transfer efficiency to the human liver.^12,20,21^

**Figure 1.**
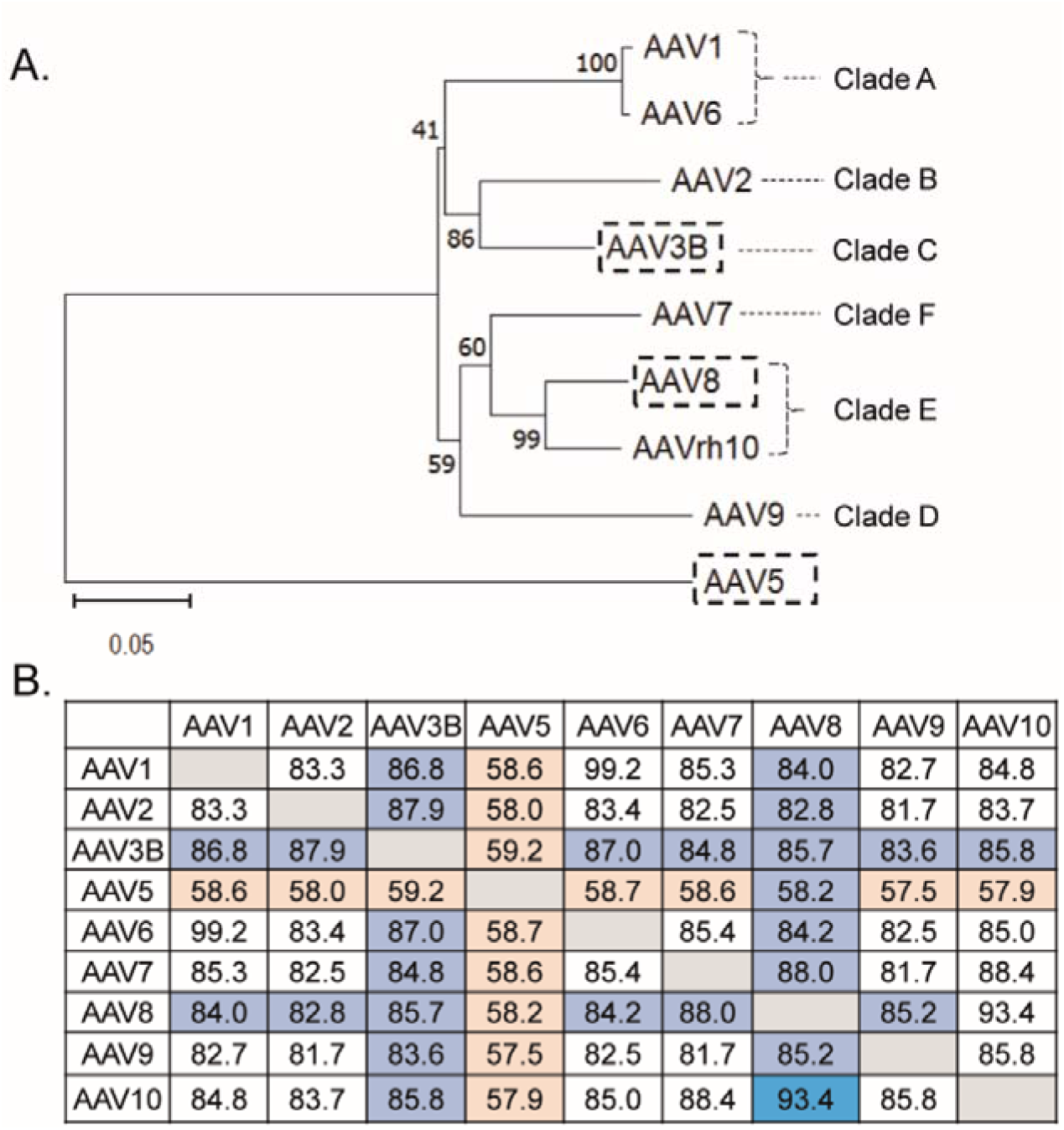
The topology based on the similarity and identity of the capsid sequences of various AAV serotypes. Phylogenetic relationships among AAV serotypes. (A) The topology of AAVs is segregated based on similarities of capsid sequences in various AAV serotypes. (B) Amino acid identities (%) are presented by pairwise alignment. Dark blue indicated >90% identity; light blue highlights 80%–90% identity and light-yellow highlights <60% identity.

### Comparison of the immunogenicity between different transgenes

First, we assessed the transduction efficacy of AAV3B, AAV5, and AAV8 vectors harboring hFIX or secNanoLuc. We injected each AAV vector, intravenously, at the same dose (0.5 × 10^12^ vg/kg) in C57Bl/6J mice, and transduction efficacy was determined by AAV genome in the liver. AAV8 vector enabled the transduction of AAV in the mouse liver when compared with AAV3B and AAV5 vectors (Figure 2A). AAV vectors harboring NanoLuc showed higher transduction efficacy than those expressing FIX in AAV3B and AAV5 (Figure 2A). However, AAV vector-mediated induction of serotype-specific NAb was almost identical between transgenes (Figure 2B). Interestingly, AAV5 induced a much higher serotype-specific NAb than those induced by both AAV3B and AAV8 capsids (Figure 2B).

**Figure 2.**
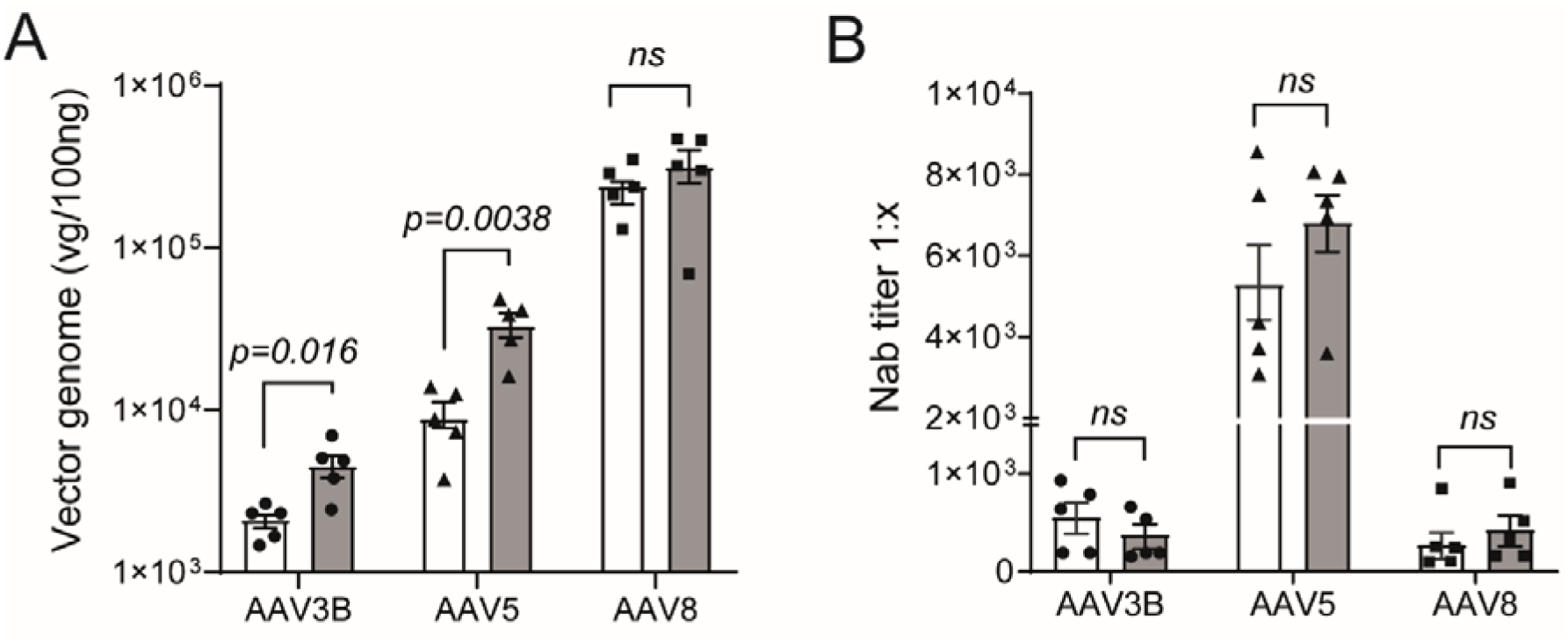
Transduction efficacy and immunogenicity of AAV3B, AAV5, and AAV8 vectors encoded human FIX or secNanoLuc in mice. AAV3B, AAV5, or AAV8 vectors harboring human FIX or secNanoLuc were intravenously injected into mice (0.5 × 10^12^ vg/kg). The serum and liver tissue were harvested 30 days after vector injection. (A) AAV vector genome in the liver was evaluated by qPCR. (B) The titers of serotype-specific NAb after the vector injection were determined using *in vitro* cell-based NAb assay. Data are presented as mean ± SEM (n = 5). Statistical analysis between two groups was performed by Student *t-*test. Actual *p* values were described in the Figure. White bar, mice treated with AAV vector harboring human FIX; gray bar, mice treated with AAV vector harboring secNanoLuc. AAV, adeno-associated virus; secNanoLuc, secreted NanoLuc; NAbs, neutralizing antibodies; ns., not significant.

Subsequently, we examined the cross-reactivity of each AAV vector-induced NAbs to other AAV serotypes (Table 1). We did not detect pre-existing NAb against AAV3B and AAV8 in mice; however, the serum samples obtained from some mice indicated the existence of a low titer of NAbs against AAV5, as described previously^22^. AAV3B vector-immunized sera showed some partial inhibition to several AAV serotypes, including AAV8 (one of 10 mice) (Table 1). We detected NAb against AAV5 in two of five mice treated with AAV3B, but these positive mice had AAV5 NAbs even before the administration. AAV5 vector-injected mice did not develop NAbs cross-reactivity against other AAVs except only partial cross-reaction with AAV1 (Table 1). On the other hand, AAV8-immunized mice developed NAbs cross-reactivity against both AAV9 and AAVrh10 serotypes, but not against either AAV3B or AAV5 serotypes (Table 1).

**Table 1.** Cross-reaction of AAV injection induced NAbs with different AAV serotypes.

| NAbs against AAV capsids |  |  |  |  |  |  |  |  |  |
| --- | --- | --- | --- | --- | --- | --- | --- | --- | --- |
| Vector injection | AAV1 | AAV2 | AAV3B | AAV5 | AAV6 | AAV7 | AAV8 | AAV9 | AAVrh10 |
| AAV3B-hFIX | 4/5 | 3/5 | 5/5 | 2/5* | 2/5 | 0/5 | 1/5 | 0/5 | 0/5 |
| AAV3B-secNanoLuc | 2/5 | 2/5 | 5/5 | 2/5* | 0/5 | 0/5 | 0/5 | 0/5 | 0/5 |
| AAV5-hFIX | 3/5 | 0/5 | 0/5 | 5/5 | 0/5 | 0/5 | 0/5 | 0/5 | 0/5 |
| AAV5-secNanoLuc | 1/5 | 0/5 | 0/5 | 5/5 | 0/5 | 0/5 | 0/5 | 0/5 | 0/5 |
| AAV8-hFIX | 0/5 | 0/5 | 0/5 | 0/5 | 0/5 | 0/5 | 5/5 | 5/5 | 5/5 |
| AAV8-secNanoLuc | 2/5 | 2/5 | 0/5 | 2/5 | 1/5 | 0/5 | 5/5 | 5/5 | 5/5 |

*n/n* = number of NAb-positive mice/mice number in each group
AAV, adeno-associated virus; secNanoLuc, secreting type of NanoLuc; hFIX, human factor 9.
\*Pre-existing NAb against AAV5 was detected in two mice and did not change after the injection.

### Re-administration of different AAV serotypes

To evaluate the possibility of re-administration of AAV vectors, C57BL/6J mice were first injected with AAV3B, AAV5, and AAV8 vectors encoding hFIX or PBS at day 0, and were then re-administered with AAV3B, AAV5, and AAV8 vectors harboring secNanoLuc at day 30. The success of the second AAV transduction was assessed by serum NanoLuc activity and AAV genome in liver at 60 days after the first administration (Figure 3). As expected from the emergence of serotype-specific NAbs after AAV vector administration, re-administration of the same AAV serotype failed to transduce in all the three groups (Figure 3). Conversely, the re-administration of different AAV serotypes were successful (Figure 3). A partial inhibition of the second transduction with AAV3B and AAV8 vectors was observed in some mice that were first treated with AAV8 and AAV3B, respectively, but the differences were not statistically significant (*p=0.45*) (Figure 3). Of note, the re-administration of AAV5 was completely achieved in mice that were previously treated with AAV3B or AAV8. Further, the transduction by re-administration of AAV3B and AAV8 succeeded in all mice first treated with AAV5 (Figure 3E and F).

**Figure 3.**
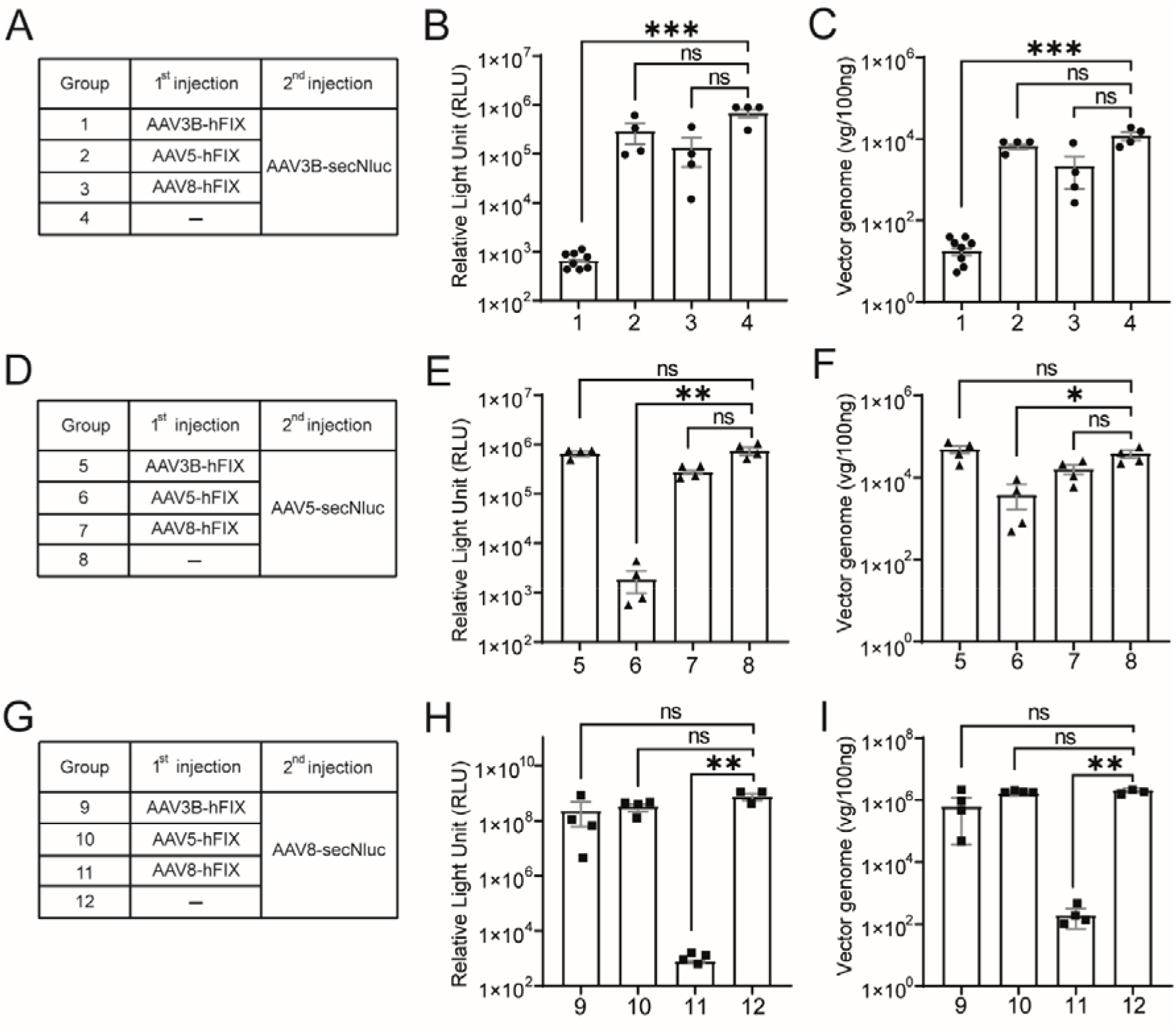
Re-administration of AAV vectors in mice. Mice were intravenously administered saline control or AAV3B, AAV5, or AAV8 harboring human coagulation factor IX (AAV-hFIX), and then received AAV3B, AAV5, or AAV8 harboring secNanoLuc (AAV-secNluc) at day 30. The sera and liver tissues were harvested 60 days after the first vector injection. (A, D, and G). The table summarizes the AAV vectors administered to each mouse. (B, E, and H) The expression level of NanoLuc in the serum was assessed by luminescence and expressed as RLU. (C, F, and I) AAV vector genome of second injection in the liver was evaluated by qPCR. Data are presented as mean ± SEM (n = 3–8). Statistical difference was analyzed by one-way ANOVA with Kruskal Wallis multiple comparisons test. \**p* < 0.05, \*\**p* < 0.01, and \*\*\**p* < 0.001. AAV, adeno-associated virus; ns., not significant.

### The changes in the titer and specificity of neutralizing antibodies after the second challenge

Next, we examined the changes in specificity and titer of NAbs after the repeated administration of the same and switching AAV serotypes (Figure 4). As shown in Figure 2, the induction of NAb by AAV5 vector was highest after the first injection (6053.5 ± 2210.1) as compared with AAV3B (427.1 ± 197.0), or AAV8 (33.7 ± 10.2) at day 30 (Figure 3). The second administration of the same AAV vectors boosted the serotype-specific NAb titer significantly (*p<0.001*), but not with the different serotypes (Figure 4).

**Figure 4.**
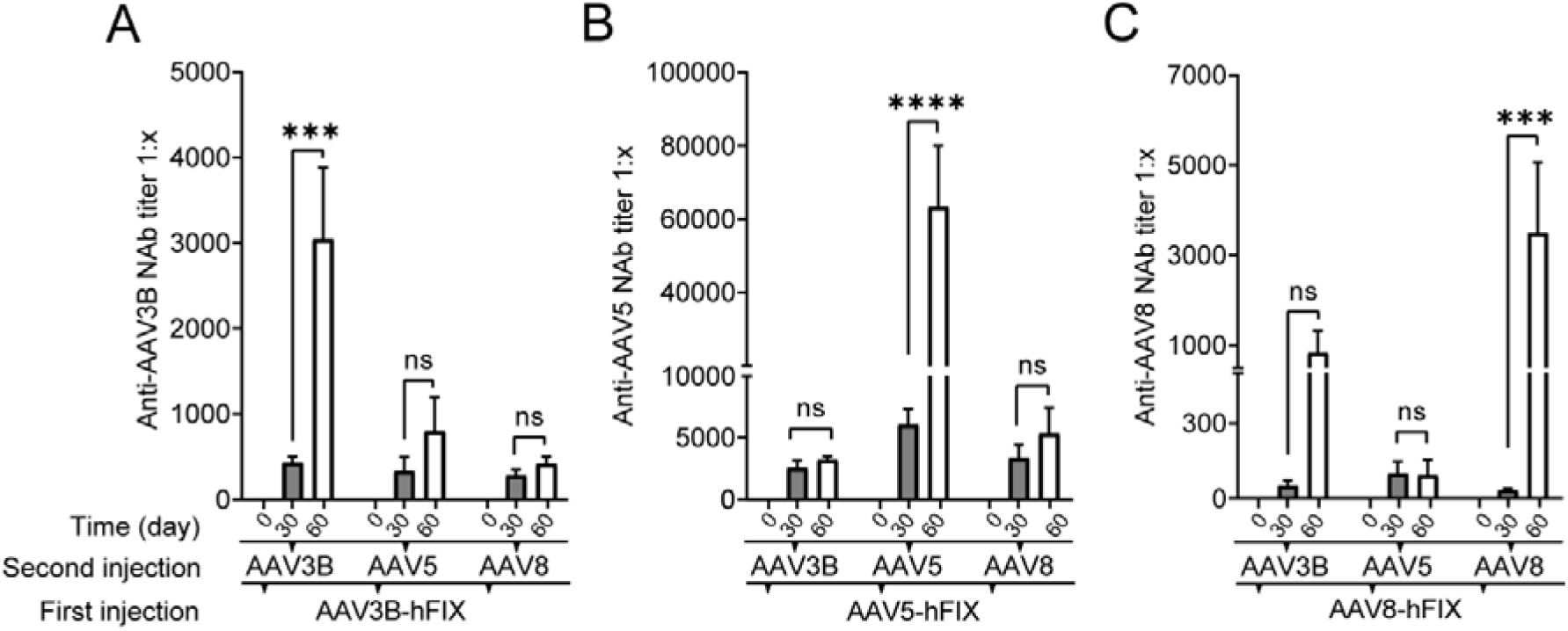
Dynamics of serotype-specific NAbs after re-administrating by switching AAV capsids. Mice were intravenously administered AAV3B (A), AAV5 (B), or AAV8 (C) harboring human coagulation factor IX. Subsequently, they received each of three different AAV capsid (AAV3B, AAV5, or AAV8) vectors harboring secNanoLuc at day 30. Sera were harvested at day 0, and 30 and 60 days after the first vector injection and NAbs against AAV3B (A), AAV5 (B), and AAV8 (C) were determined. Data are represented as mean ± SEM (n = 4–8). Statistical difference was analyzed by two-way ANOVA with Turkey’s multiple comparisons test. *p < 0.05, **p < 0.01, ***p < 0.001 and ****p < 0.0001. AAV, adeno-associated virus; ns., not significant.

Furthermore, we evaluated the cross-reactivity of the sera after two administrations of AAV vectors against the various AAV serotypes (Table 2). The sera obtained from two doses of AAV3B inhibited the transduction with AAV1 and AAV2 capsids, but not against AAV5 and AAV9 (Table 2). The sera from the second doses of AAV8 were cross-reacted with AAV3B, AAV5, AAV9, and AAVrh10, while being partially reactive against AAV1 and AAV2 vectors (Table 2). Interestingly, repeated administration of AAV5 tended not to induce NAbs cross-reactivity with the other AAV serotypes; only one mouse developed NAbs cross-reactivity with AAV1 and AAV8 (Table 2). Likewise, the sequential administration of either AAV3B or AAV8 did not induce NAbs cross-reactivity with AAV5 capsid. In addition, the re-administration of the different serotypes produced NAbs that reacted with a wide range of AAV serotypes (Table 2). However, NAbs against AAV7 were generated only in the repeated administration of AAV3B, while the other combinations of re-administration did not (Table 2).

**Table 2.** Cross-reaction of repeated AAV injection via same serotype induced NAb with different AAV serotypes.

|  |  | NAbs against AAV capsids |  |  |  |  |  |  |  |  |
| --- | --- | --- | --- | --- | --- | --- | --- | --- | --- | --- |
| First injection | Second injection | AAV1 | AAV2 | AAV3B | AAV5 | AAV6 | AAV7 | AAV8 | AAV9 | AAVrh10 |
| AAV3B-hFIX | AAV3B-secNanoLuc | 8/8 | 8/8 | 8/8 | 2/8* | 4/8 | 3/8 | 3/8 | 0/4 | 2/8 |
| AAV3B-hFIX | AAV5-secNanoLuc | 1/4 | 1/4 | 4/4 | 4/4 | 0/4 | 0/4 | 1/4 | 0/4 | 1/4 |
| AAV3B-hFIX | AAV8-secNanoLuc | 2/4 | 4/4 | 4/4 | 0/4 | 1/4 | 0/4 | 4/4 | 4/4 | 4/4 |
| AAV5-hFIX | AAV3B-secNanoLuc | 1/4 | 2/4 | 4/4 | 4/4 | 0/4 | 0/4 | 1/4 | 0/4 | 0/4 |
| AAV5-hFIX | AAV5-secNanoLuc | 1/4 | 0/4 | 0/4 | 4/4 | 0/4 | 0/4 | 1/4 | 0/4 | 0/4 |
| AAV5-hFIX | AAV8-secNanoLuc | 0/4 | 3/4 | 4/4 | 4/4 | 0/4 | 0/4 | 4/4 | 0/4 | 4/4 |
| AAV8-hFIX | AAV3B-secNanoLuc | 3/4 | 4/4 | 4/4 | 0/4 | 2/4 | 0/4 | 4/4 | 3/4 | 4/4 |
| AAV8-hFIX | AAV5-secNanoLuc | 3/4 | 0/4 | 3/4 | 4/4 | 0/4 | 0/4 | 4/4 | 3/4 | 4/4 |
| AAV8-hFIX | AAV8-secNanoLuc | 1/4 | 3/4 | 4/4 | 4/4 | 0/4 | 0/4 | 4/4 | 4/4 | 4/4 |

*n/n* = number of NAb-positive mice/mice number in each group
AAV, adeno-associated virus; secNanoLuc, secreting type of NanoLuc; hFIX, human factor 9.
\*Pre-existing NAb against AAV5 was detected in two mice and did not change after the injection.

## Discussion

Generally, AAV vector-based gene therapy is capable of long-term gene expression to treat a variety of genetic disorders. Specifically, the primary application of AAV vectors is for the expression of therapeutic genes in the central nervous system or liver. While neural cells of the central nervous system do not divide, it is possible that the therapeutic effects of AAV-mediated gene therapy in the liver gradually diminish over time with the proliferation of hepatocytes. Although re-administration is one possible solution to recover the therapeutic level; however, the second challenge of the immune system with the same AAV vector capsid is impossible due to the high titer of NAb induced with the previous administration. Hence, various strategies for re-administration are currently being developed. Repeated administration of the same AAV vector has been successful with several immunomodulators in the liver with AAVs in murine and monkeys.^23,24^ Recently, degrading the circulating IgG by endopeptidase liver is being tested to remove pre-existing NAbs; thus, enabling the infusion of AAV vectors in mice and monkeys.^25^ In the present study, we demonstrated the successful re-administration of liver-targeting serotypes of AAV3B, AAV5, and AAV8 in mice by switching vector serotypes having confirmed the absence of cross-reactivities of NAb after the first administration.

The success of re-administration by another serotype depends on the specificity of the NAbs produced after the administration of the AAV vectors. Upon the infusion of the vectors, mice and non-human primates developed high titers of anti-AAV neutralizing antibodies, and it is also the case for humans. It was reported that the treatment-induced NAb is relatively serotype-specific in animal models.^26,27^ The experiments with canine hemophilia model treated with AAV2, AAV6, and AAV8 demonstrated limited cross-reactivity to other capsids.^28^ In addition, the treatment with AAV5 vectors in murine and monkeys did not induce NAbs which are cross-reacted with AAV1.^16^ Several clinical trials with AAV vectors are currently underway; hence, we could not obtain sufficient data about seroreactivity of NAbs induced by AAV injection in human beings. Systemic AAV vector (AAV2) administration reportedly elicited NAbs cross-reacted with multiple serotypes, and they persisted for a long time in a 15-year follow-up. However, we could not assess whether NAb was induced by AAV vector or not, because pre-existing NAb was not addressed in this study.^8^ In the current clinical studies, it is necessary to clarify the serotype specificity of NAbs developed after the administration of the vectors.

In human beings, the NAb-producing mechanism might be different in natural AAV infection because the pre-existed of NAbs in the human sample without AAV treatment could cross-react with multiple AAV serotypes. The seroprevalence of NAbs against AAV due to natural infection in humans varies from 20 to 80% of the parent population.^29^ We have recently examined the seroprevalence of NAbs against nine AAV serotypes and found most NAbs reacted with multiple AAV serotypes in Japanese.^15^ AAV infection and NAb are also found in animals such as rodents, cats, pigs, dogs, horses, and chimpanzee.^27,30–32^ Although serotypes and NAb titer vary depending on animals’ species, cross-reactivity to many serotypes is common, as in the case with humans. A possible interpretation of this might be related to repeated exposure to the living environment and latent AAVs, integrated into the chromosome and re-activated with the infection of helper virus, including adenovirus.^14,33^ Indeed, our data suggest that the second challenge with AAV induced high titer of NAbs which cross-reacted with many AAV serotypes.

The specificity of NAbs induced by the treatment with AAV5 vector seemed to be different from those induced by AAV3B and AAV8. The first injection of AAV5 induced a very high titer of serotype-specific NAbs in mice and further increased the NAb titer after the second administration. Even this extremely high NAb titer did not cross-react with other serotypes. On the contrary, the high titer of NAbs induced by the second administration of AAV3B and AAV8 abolished the transduction with multiple AAV serotypes. This characteristic feature is probably due to the capsid sequence similarity of AAV5 when compared to the other AAVs. Specifically, AAV5 is the most distant serotype based on topology.^34,35^ In addition, there are interesting reports on NAbs for AAV5. For instance, AAV5 vector-mediated hFIX gene transfer was reportedly achieved with the presence of pre-existing NAbs, albeit a definitive explanation remains to be established.^22,36^ One possible explanation might be the dose of AAV5 employed in the previous study. High-dose of the vector is required for the efficient transduction with AAV5 because of its low transduction efficacy to liver cells.^37,38^ We previously showed that a high-dose vector (1 × 10^13^ vg/kg) could evade NAbs for liver transduction with AAV8 vector.^39^ confirmed transduction with a lower vector dose (0.5 × 10^12^ vg/kg) was inhibited by 1:1 Nab ratio even with an AAV5 vector (data not shown). Although NAbs induced by the first AAV5 abolished re-administration in this study, further investigation is required to determine whether the AAV5 capsid itself has the property to evade NAbs.

This study has several limitations. Here, we only assessed humoral immune response against the AAV vector for a short period in a mice model. Further, we did not evaluate the durability of transgene and the kinetics of NAb titer during the time. We should collect information about not only humoral immunity but also innate/cellular immunity to comprehensively understand vector immunogenicity. In addition, it is necessary to carefully examine the immune response to vectors in ongoing human clinical trials, as well as in preclinical studies in non-human primates over a long period.

In summary, we validated the possibility of re-administering AAV vectors for liver-directed gene therapy to change the serotypes in mice. The use of AAV vectors has been an effective treatment strategy against hemophilia and other genetic diseases because of its long-term gene expression in the liver. However, it is not known how long the therapeutic effect will be maintained, *e.g*., 10 years or longer, and the therapeutic efficacy varies among individuals. In this context, to further facilitate the effectiveness of AAV vector therapy, it is also necessary to prepare several serotype formulations for re-administration in treating one disease.

## Acknowledgments

This work was supported by AMED [JP18pc0101030]. The BD LSR-Fortessa flow cytometer, Optima XE-90 was subsidized by JKA through its promotion funds from KEIRIN RACE. We thank Yaeko Suto, Mika Kishimoto, Tamaki Aoki, Sachiyo Kamimura, Mai Hayashi, Yuiko Ogiwara, Nagako Sekiya, Noguchi Tomoko, Hiromi Ozaki, and Hiroko Hayakawa of Jichi Medical University for their technical assistance. The graphical abstract was created using BioRender.com.

## Author contributions

N.B. designed the study, performed the experiments, analyzed the data, and wrote the manuscript. Y.K. designed the study, performed the experiments, and revised the manuscript. M.H., N.K., T.H., and H.M. designed the study, analyzed the data, and revised the manuscript. T.O. designed the study, analyzed the data, and wrote the manuscript. All authors approved the final version of the manuscript.

## Data availability statement

The data that support the findings of this study are available from the corresponding author upon reasonable request.

